# scLASER: a robust framework for simulating and detecting time-dependent single-cell dynamics in longitudinal studies

**DOI:** 10.64898/2026.05.04.722712

**Authors:** Lauren A Vanderlinden, Juan Vargas, Jun Inamo, Jade Young, Chuangqi Wang, Fan Zhang

**Affiliations:** Department of Biomedical Informatics, University of Colorado School of Medicine, Aurora, CO, USA; Department of Medicine Rheumatology, University of Colorado School of Medicine, Aurora, CO, USA; Laboratory for Human Immunogenetics, RIKEN Center for Integrative Medical Sciences, Yokohama, Japan; Department of Microbiology and Immunology, Keio University School of Medicine, Tokyo, Japan; Department of Immunology and Microbiology, University of Colorado School of Medicine, Aurora, CO, USA

**Author notes:** **Corresponding author:** Fan Zhang.

## Abstract

Longitudinal single-cell clinical studies enable tracking within-individual cellular dynamics, but methods for modeling temporal phenotypic changes and estimating power remain limited. We present scLASER, a framework detecting time-dependent cellular neighborhood dynamics and simulating longitudinal single-cell datasets for power estimation. Across benchmark experiments, scLASER shows consistently higher sensitivity than traditional cluster--based approaches, with particularly pronounced gains in rare cell types and non-linear temporal patterns. Applications to inflammatory bowel disease (95,813 cells, 38 patients) reveal treatment-responsive NOTCH3+ stromal trajectories with high cell type discrimination (AUC > 0.92), while analysis of COVID-19 data (188,181 cells, 84 patients) identifies three distinct axes of T cell activity (cytotoxic effector, NK immunoreceptor signaling, and interferon-stimulated gene programs) over disease progression. scLASER enables robust longitudinal single-cell analysis and optimization of study design.

**Teaser:** A new framework detects time-dependent cellular dynamics and enables power estimation for longitudinal clinical single-cell studies.

## Introduction

Longitudinal studies using large single-cell multi-omics allow the tracking of within-individual changes in cellular states over time. This enables stronger causal inference about treatment response, disease progression, and immune state transitions, rather than inferring dynamics from cross-sectional snapshots^1^. Longitudinal approaches based on variance decomposition^2^ applied to bulk or single-cell gene expression data^3^ focus primarily on identifying stable versus variable features rather than modeling directional temporal trajectories. Critically, analyzing individual cell types in isolation may miss coordinated changes in the cellular microenvironment where multiple cell types shape clinical outcomes. Methods for robustly modeling how these multicellular compositions evolve over time in response to treatment or disease progression remain limited, restricting analyses to cross-sections at individual time points^4,5^ In practice, such datasets are also inherently nested (cells within samples within participants), unbalanced across visits, and affected by technical artifacts and missing data, requiring specialized analytical approaches.

Another challenge in these study designs is estimating the power to detect temporal changes in cellular neighborhoods associated with clinical outcomes. Although methods exist to simulate longitudinal gene expression^6,7^, or test for differential neighborhood abundance^8^, none provide simulation frameworks for power estimation in longitudinal clinical studies. Consequently, researchers lack a principled framework to assess power or optimize study design before costly data collection.

To address these limitations, we develop scLASER (Longitudinal Analysis with Simulator and Effect detectoR), a unified framework for detecting time-dependent cellular neighborhood dynamics and simulating realistic longitudinal datasets for power estimation. scLASER integrates raw single-cell profiles with metadata to construct neighborhoods, followed by unsupervised variance decomposition or covariance-based supervised learning to generate embeddings, to identify temporal trajectories contributing to clinical outcomes like treatment response (**Figure 1**).

**Figure 1.**
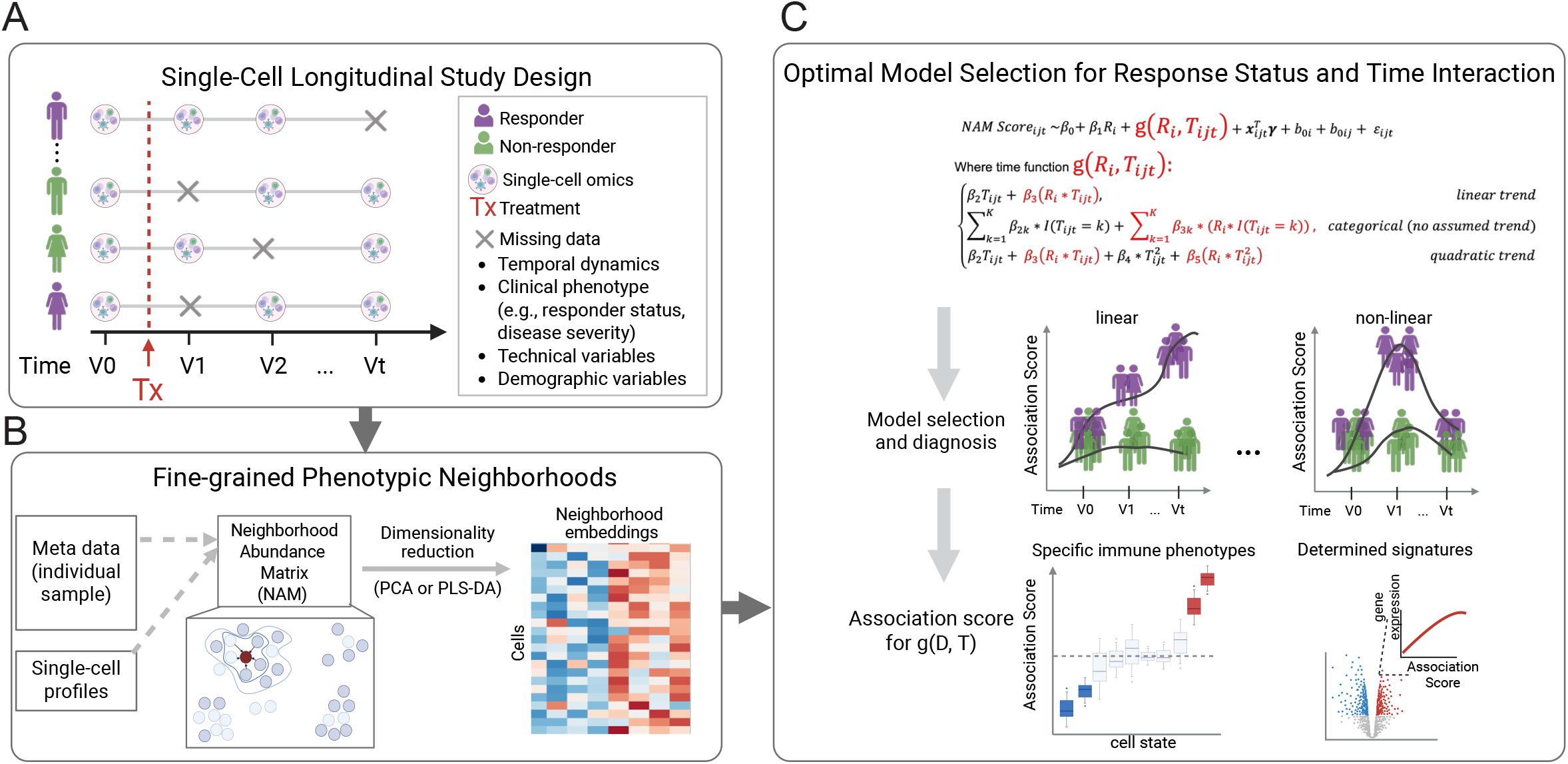
scLASER analysis workflow. Overview of scLASER pipeline for **A.** longitudinal single-cell studies, including **B**. phenotypic neighborhoods construction, **C**. model selection for clinical outcome-by-time interactions, and association score interpretation.

It generates realistic longitudinal datasets with user-defined variables, including cell type neighborhood dynamics, clinical effect sizes, linear and nonlinear temporal patterns, technical variation, and demographic factors, thereby enabling power calculations and design optimization (**Figure 2A**). We validated scLASER through systematic simulations of increasing complexity, benchmarking against cluster-based frequency approaches. We demonstrated its ability to reveal rare and novel cell phenotypes, including a treatment-responsive stromal population and orthogonal lymphocyte activation axes linked to infectious disease severity.

**Figure 2.**
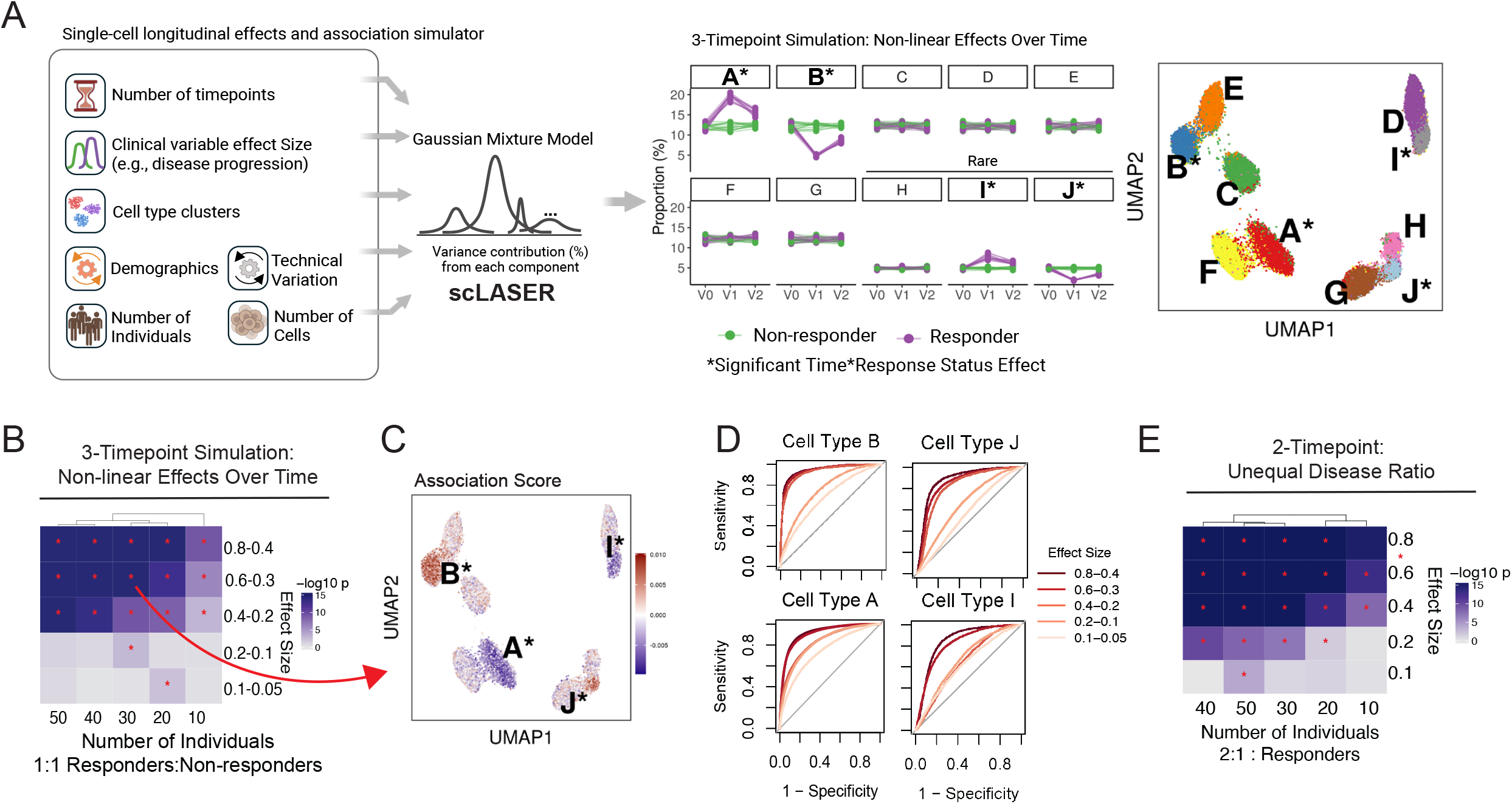
scLASER simulation-based validation. **A.** Simulation framework of scLASER; example three-timepoint simulations show responders versus non-responders trajectories (V0-V2) and corresponding UMAP. **B**. Detection performance in three-timepoint simulations across varying numbers of individuals and effect sizes with heatmaps indicating LRT p-values. * statistical significance. **C**. UMAP of simulated cell types colored by scLASER association scores, * indicates cell types A and B (major) and I and J (rare) are significant time×response status effects. **D**. ROC curves demonstrating classification performance for cell types with simulated time×response status effect. **E**. Two-timepoint simulation and performance with unbalanced 2:1 non-responders:responders ratio.

## Results

### Simulation-based validation demonstrates scLASER’s high sensitivity across effect sizes

We first evaluated scLASER’s performance through simulations of longitudinal datasets with responders to treatment and non-responders. Simulations included 10 cell types, with four carrying response-by-time signals (cell types A, B, I, and J). In two-visit scenarios, we simulated effect sizes from 0.1 to 0.8, representing both expansions and depletions in responders vs. non-responders. scLASER’s leading association score, captured by neighborhood abundance matrix (NAM) PC1 (**Methods**), clearly separated groups at moderate to large effect sizes (0.6–0.8) and detected weaker signals (effect size 0.1). Receiver operating characteristic (ROC) analysis demonstrated discrimination for true positives, cell types A and I (expanded), and B and J (depleted), in responders with area under the curve (AUC) exceeding 0.9 for effect sizes above 0.4, and moderate sensitivity (AUC 0.7–0.8) at small effects. Cell types without imposed signals showed AUC near 0.5 across all scenarios, confirming specificity (**Supplementary Figure 1**).

### scLASER outperforms cluster-based frequency approaches for rare cell types and non-linear patterns

To benchmark scLASER’s sensitivity and granularity, we implemented a cell-type cluster-based frequency approach incorporating time and response status interaction effect to test performance (**Methods**). Across 200 randomly generated datasets with effect size 0.6, 30 individuals, and 100 cells per major type (**Figure 2**), scLASER achieved higher sensitivity than the cluster-based test (97.5% vs 94.4%), and most pronounced gains for rare types (96.7% vs 88.8%) (**Supplementary Figure 4**). In three-visit simulations (V0, V1, V2), we generated both linear trends (monotonic expansion/depletion) and non-linear bump patterns (signal peaks at V1). Model selection via AIC correctly identified linear time parameterization for linear trends and quadratic or categorical time for nonlinear patterns, validating the adaptive model-selection strategy (**Methods**; **Supplementary Figures 2,3**). Across 200 simulated datasets with the non-linear effect size of 0.2 at V1 and then decreasing to 0.1 at V2 (**Figure 2B**), scLASER has a sensitivity of 87.8% while the frequency-based method had a sensitivity of 30.9%. Again, gains are more pronounced specifically in rare cell types (sensitivities of 87.1% vs 4.5% respectively) (**Supplementary Figure 4**). ROC analysis demonstrated consistent scLASER classification performance across different non-linear effect patterns, with AUCs exceeding 0.8 for effect sizes ranging from 0.4 at V1 to 0.2 at V2 (**Figure 2C**). Simulation with a 2:1 ratio of non-responders:responders resulted in slightly reduced sensitivity as expected (**Figure 2D**).

### Supervised approach captures time-dependent neighborhood signals in high-variance datasets

PCA identifies maximal variance but may miss clinically relevant, time-dependent outcome signals that contribute less to total variability. In contrast, partial least squares discriminant analysis (PLS-DA)^9^ directly leverages covariance with the outcome, potentially increasing sensitivity to structured effects. To illustrate this, we simulated a higher-variance dataset with one cell type showing a time-dependent response effect and another cell type showing only a main response effect independent of time (**Figure 3A**). Comparing PCA, binary PLS-DA (responder vs non-responder) and six-level PLS-DA (responder status at each timepoint) revealed two key findings. First, PCA (PC4) and six-level PLS-DA (component 1) showed comparable sensitivity to time-dependent effects, both correctly identifying the interaction cell type (**Figure 3B**). The six-level PLS-DA showed slightly improved specificity as it was less influenced by the main-effect-only cell type. Second, the binary PLS-DA failed to detect the time-dependent pattern, instead capturing only the main effect, highlighting that outcome specification critically impacts signal detection. Differences between PCA and PLS-DA loadings reflect their different objectives (**Supplementary Figure 5**). For datasets with high compositional variance, PLS-DA using a composite outcome between time and clinical outcome is recommended as it can appropriately model the covariance between clinical outcomes and cellular neighborhoods. scLASER enables users the flexibility to apply either technique depending on analytical goals.

**Figure 3.**
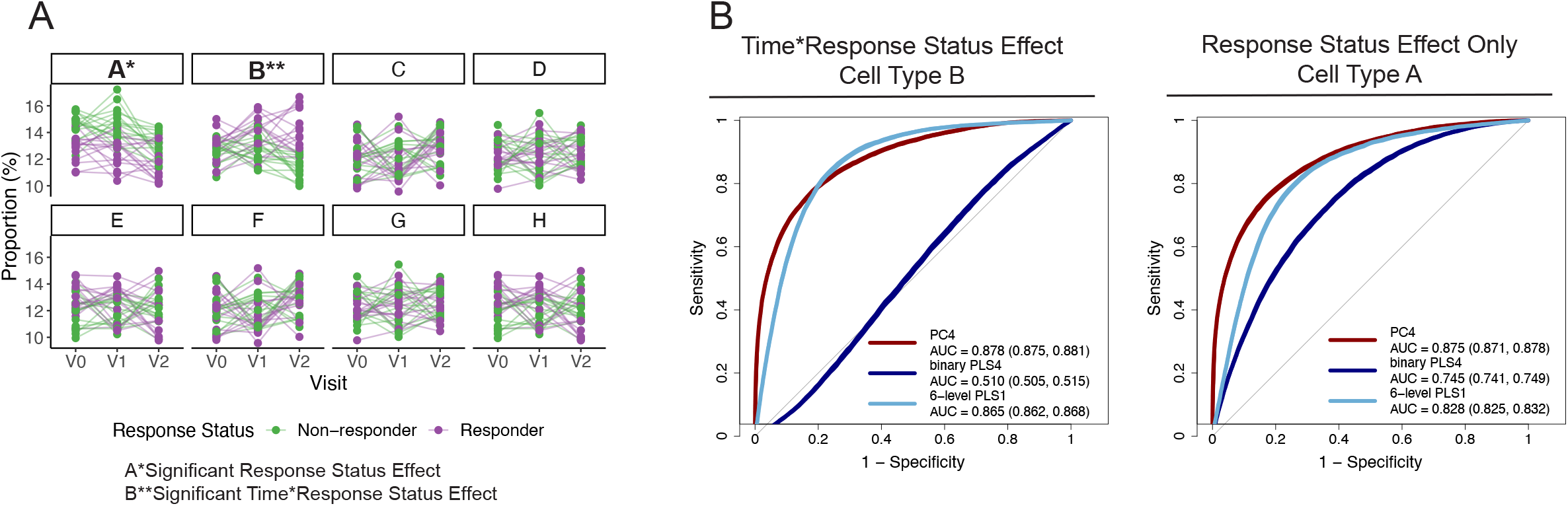
Sensitivity of dimensionality reduction methods to longitudinal response-associated cell type dynamics. **A.** Cell type proportion trajectories across three visits in high variability simulatwion, * indicates a main response effect and ** indicates significant time×response status effect. **B**. ROC comparison of dimensionality reduction approaches for detecting time×response status effects (left) versus response-only effects (right).

### scLASER identifies treatment-responsive NOTCH3+ stromal populations in IBD

To apply scLASER on real longitudinal data with cell-type-specific, time-dependent effects, we analyzed stromal cells (95,813 cells) from patients with inflammatory bowel disease (IBD)^4^ (38 participants: 22 ulcerative colitis, 16 Crohn’s disease; 192 samples: 98 UC, 94 CD) collected across five biopsy sites, including paired pre- and post-adalimumab treatment samples (**Figure 4A**). The identified time-by-remission association score increased post-treatment in remission patients only (**Figure 4B**, LRT p=3e-4), indicating their depletion correlated with treatment response. Visualizing the association score confirmed that this axis captured the segregation of *NOTCH3+* stromal populations, annotated based on original study (**Figure 4C-E**). Predictive modeling using abundances of three *NOTCH3+* populations achieved strong discrimination of remission status (AUC=0.953, 0.950, and 0.926, respectively; **Figure 4F, Supplementary Figure 6**). Genes correlated with the association score included key markers of stromal activation and extracellular matrix remodeling, including *NOTCH3, ACTA2, MEF2C*, and *RGS5* (**Figure 4G-H**). Notably, this longitudinal signal was identified by modeling unmatched biopsy sites and mixed UC/CD subtypes within a unified framework, enhancing the power to uncover patterns missed by the original non-longitudinal analyses. These findings improve upon emerging evidence that intestinal fibroblast and pericyte states mediate treatment response in IBD^10^.

**Figure 4.**
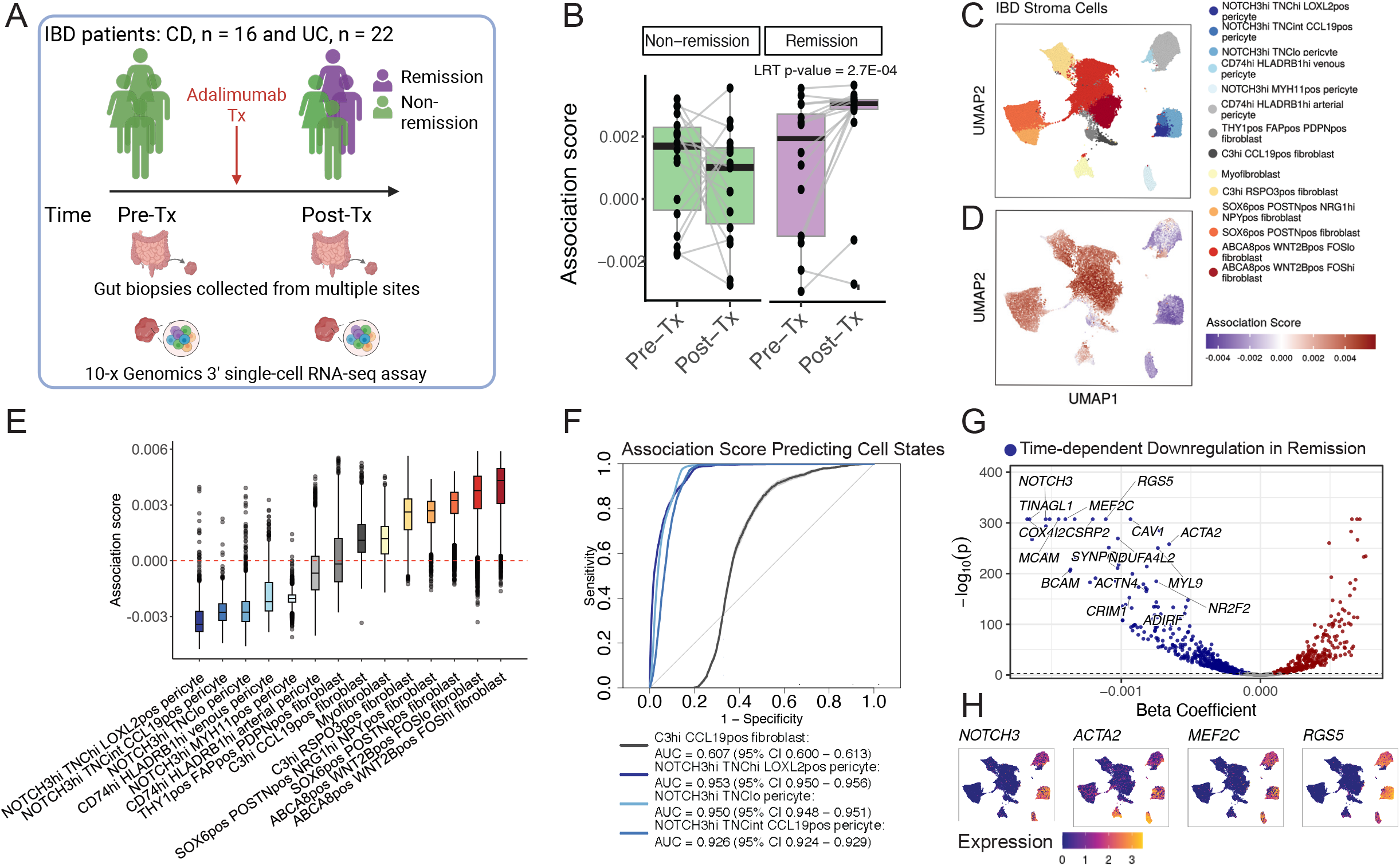
scLASER identifies treatment-responsive stromal neighborhoods in inflammatory bowel disease. **A.** IBD study design: 38 patients with paired pre- and post-adalimumab biopsies from multiple gut sites, stratified by remission status. **B**. Association score showing a time×remission interaction (LRT p=2.7e-4). **C**. Stromal cell UMAP colored by cell type and **D**. scLASER association score revealing treatment-responsive neighborhoods. **E**. Association score distributions across stromal subsets. **F**. ROC curve demonstrating the association score predictive performance for NOTCH3+ cluster abundance. **G**. Genes correlated with the association score and **H**. UMAP expression of stromal activation markers.

### Three distinct T cell activation axes emerge during COVID-19 progression using scLASER

To assess scLASER at increased temporal resolution and unbalanced sampling, we analyzed a COVID-19 cohort tracking participants across up to four time points from symptom onset, with a focus on identifying time-by-progression interactions in T and NK cells (188,181 cells, 84 patients, 19 progressors, 65 non-progressors)^5^ (**Figure 5A-B**). With scLASER, we identified three activation axes associated with temporal abundance shifts between progressors and non-progressors. NAM PC1 is an inflammatory effector axis (LRT p-value = 7e-04, **Figure 5C-D**) capturing a cytotoxic T cell program marked by granule-mediated killing (e.g., *NKG7, PRF1, GZMB, GNLY, CCL5*), antigen presentation (e.g., *HLA-A/B, HLA-DR, CD74*), and type I interferon signaling, as confirmed by gene set enrichment (**Figure 5E-F**). Next, axis 2, NAM PC2 (**Figure 5G-H**, LRT p-value = 1e-04), represented an upstream NK immunoreceptor/ITAM–SYK signaling program dominated by FcεR/ITAM adapters (e.g., *FCER1G, TYROBP/DAP12*), proximal kinases (e.g., *SYK, LYN, PLCG2, LAT2*), and NK identity markers (e.g., *KLRD1/CD94, KLRF1/NKG2D, NCR1*), enriched for signal transduction and vaccine-response modules (**Figure 5I-J**). Axis 3, NAM PC12 (LRT p-value = 7e-04, **Figure 5K-L**), captured dominant interferon-stimulated gene signatures (n = 13)^5^, including *IFIT1/2/3, OASL, STAT2, ISG15* (**Figure 5M-N**). This pattern aligns with reports that severe COVID-19 features an early IFN surge followed by immune exhaustion, whereas controlled infections sustain balanced antiviral responses without hyperactivation. Beyond the interferon axis identified in the original study using only two time points, our comprehensive longitudinal analyses revealed additional parallel axes driven by cytotoxic T cells and NK cells, implicating previously unrecognized cell programs linked with disease severity.

**Figure 5.**
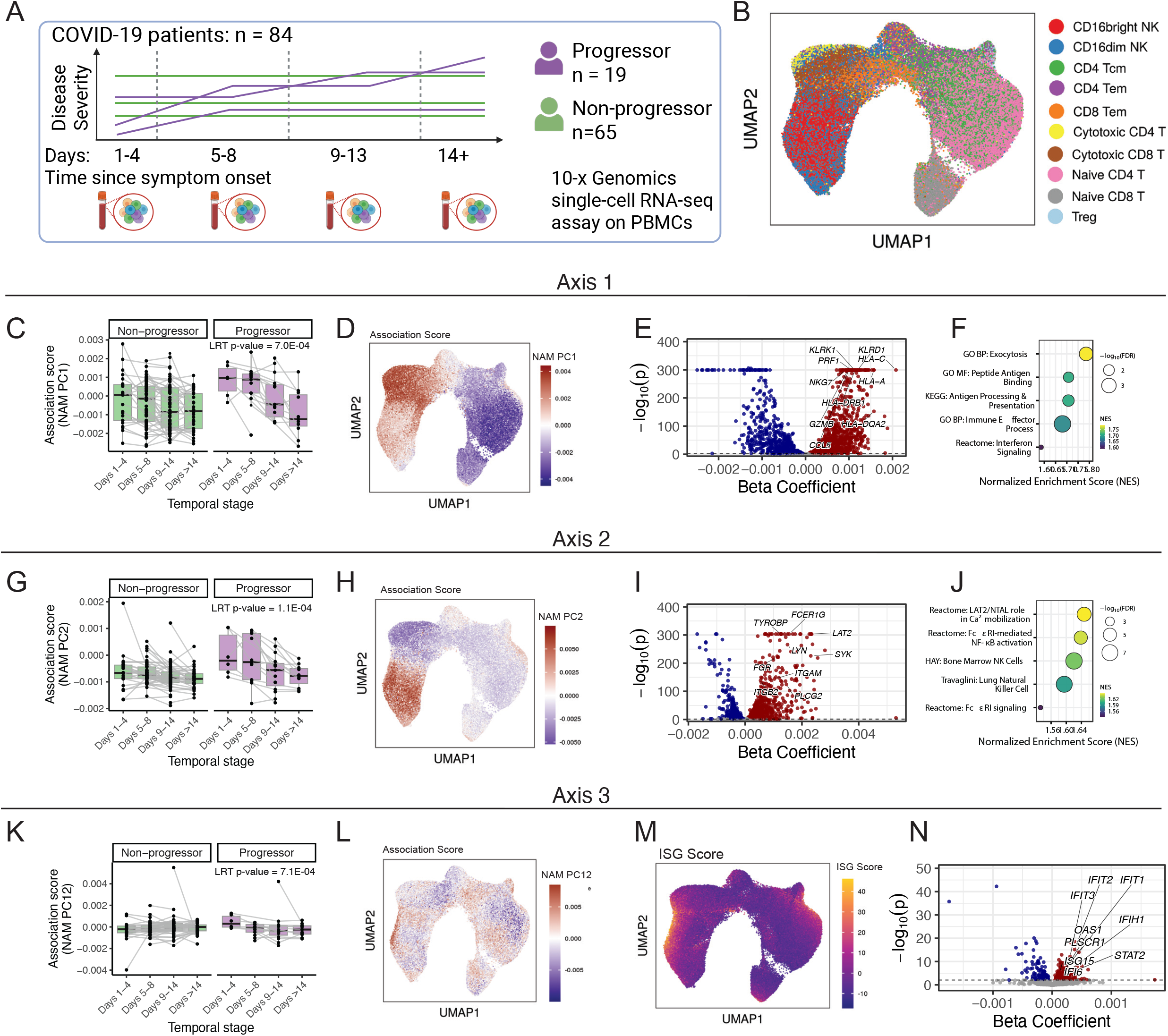
scLASER reveals progression-associated immune trajectories in COVID-19. **A.** COVID-19 study design: 84 participants across temporal stages post symptom onset. **B**. T/NK cell clusters. **C,G,K**. Longitudinal association score for three different axes: cytotoxic CD8+ effector, CD16+ NK/ADCC, and ISG-high, stratified by progression across temporal stages. **D,H,L**. UMAP colored by corresponding axis association scores. **E,I,N**. Gene associations with top markers labeled. **F,J**. Enrichment processes of axis 1 (cytotoxic granule-mediated killing) and axis 2 (Fc receptor/NK cytotoxicity). **M**. UMAP of type I interferon-stimulated genes recapitulates axis 3.

## Discussion

scLASER addresses critical gaps in longitudinal single-cell analysis by capturing time-dependent cellular dynamics and enabling power-informed study design. Our method is able to tackle two-timepoint designs as most single-cell clinical studies, like drug response studies, are based on two timepoints. scLASER can further infer the dynamics using three and higher-order timepoints as in theory three timepoints can yield dynamical systems more comprehensively^11^. While scLASER demonstrated effective identification of neighborhood-level compositional shifts often missed by frequency-based approaches, future extensions are possible. For example, while neighborhood association scores proved stable and interpretable in an unsupervised manner, supervised methods such as PLS-DA may enhance signal detection when clinical comparisons are well-defined but are more sensitive to outliers; thus, PCA is recommended by default with PLS-DA for hypothesis refinement (**Supplementary Figure 5**). The framework can be generalized to other single-cell modalities, including spatial transcriptomics. By simulating datasets under user-defined effect sizes, sampling intervals, cohort compositions, scLASER informs study design to ensure sufficient power for detecting clinically meaningful temporal signals. Overall, scLASER provides a rigorous and practical solution for modeling longitudinal dynamics, with an open-source R package available at https://github.com/fanzhanglab/scLASER.

## Materials and Methods

### scLASER Longitudinal Analytical Method

#### Data structure construction

The longitudinal analysis of scLASER (Figure 1A) begins by constructing a neighborhood abundance matrix (NAM) from batch-corrected low-dimensional embeddings and sample metadata. NAM construction follows the approach by Reshef et al^8^, using k-nearest neighbors to quantify local cell-state abundance for each sample. We focus on NAM rather than the low-dimension embeddings that go into constructing NAM because NAM captures changes in the local cellular neighborhood structure, reflecting shifts in cell-state composition and abundance across samples that are not evident from global expression variation alone. This emphasis on neighborhood-level changes allows us to directly quantify dynamic shifts in cell-state proportions between groups, which aligns with the central goal of longitudinal data analysis.

#### Dimensionality reduction

We then perform dimensionality reduction on the NAM to generate NAM scores that serve as the outcome variable in our statistical models. scLASER supports both unsupervised principal component analysis (PCA) and supervised partial least squared discriminant analysis (PLS-DA), each suited to different analytical contexts. PCA identifies axes of maximal variance in the NAM without reference to outcome variables when the goal is to characterize the dominant patterns of neighborhood variation, whereas PLS-DA is supervised and explicitly models covariance between neighborhoods and the specific outcome of interest (Eq 1). For each component h=1,…,H PLS-DA finds weight vectors (w_h_ and c_h_) that maximize the covariance between NAM and clinical outcome. The choice of outcome for PLS-DA is critical. Binary outcomes (e.g., responder vs non-responder) provide straightforward discrimination, but may discard important temporal dynamics. In longitudinal studies, specifying an outcome that preserves the time-dependent information, such as a composite term that incorporates both time and clinical phenotype (e.g., responder_V0, responder_V1, responder_V2, non-responder_V0, non-responder_V1, non-responder_V2). Because PLS-DA operates on covariance structure, subject-level effects were adjusted for in the NAM prior to PLS-DA using ComBat^12^; however, any method that adequately removes subject-specific effects can be used. This approach maintains the supervised nature of PLS-DA while modeling temporal trajectories. By default, scLASER uses PCA for creating NAM scores, as its unsupervised nature preserves flexibility for downstream modeling.

#### Analytical modeling

NAM scores are then used as the dependent variable in a mixed-effects model (LMM) framework. Traditionally, for continuous and normally distributed outcomes such as NAM scores, LMMs are considered the gold standard for longitudinal data analysis^13,14^. This framework handles the correlation between repeated measures within subjects, accommodates unbalanced data with varying number of observations per participant and irregular visit timing, is robust to missing data and allows modeling of both population-level changes and individual trajectory variation. Importantly, LMMs can also flexibly represent nested correlation structures; in our context, this corresponds to multiple cells measured within a sample and multiple samples measured within a participant. By structuring random effects accordingly, we account for correlation across levels of the data hierarchy.

We considered time primarily as a visit-based variable to capture clinically relevant study design features (e.g., baseline, follow-up, post-treatment) to allow for clinical interpretation of results. Most single-cell longitudinal studies are designed around such protocol-defined clinical timepoints, rather than irregular or opportunistic sampling, which makes visit-based modeling both practical and interpretable. We recognize that the spacing between visits may vary across participants, and therefore recommend performing a sensitivity analysis that includes the actual chronological time between visits as a covariate. This approach allows us to preserve the interpretability of discrete visits while assessing the robustness of results to heterogeneity in visit timing.

#### Model selection

Our primary focus is on modeling the interaction between time and clinical outcome of interest, for example, treatment response status (or other defined grouping variables such as progression status), as this enables us to identify changes in cellular features over time that differ between groups. Building on this framework, we model the time-response interaction, we employ three different approaches: time as a continuous variable (e.g., 0, 1, 2 and so on for baseline, follow-up 1, follow-up 2), time as a categorical variable, or time with a quadratic term (Eq. 1). The choice of time parameterization depends on the number of timepoints available. For studies with more than three timepoints, we compare all three approaches and select the model with best fit via the Akaike information criteria (AIC), while studies with only two timepoints default to categorical time modeling. All models include the corresponding main effects and lower-order terms of the interaction, along with user-specified covariates for adjustment. Random effects are structured as samples nested within participants to account for the longitudinal correlation structure. After selecting the best-fitting model based on AIC, we perform a likelihood ratio test (LRT) comparing the selected model to a nested null model that excludes the interaction term(s) but retains all main effects and covariates. This LRT provides a formal statistical test of whether the time-response interaction is significantly associated with the NAM score. NAM scores with a LRT p-value < 0.05 are considered significantly associated with the time-response interaction.

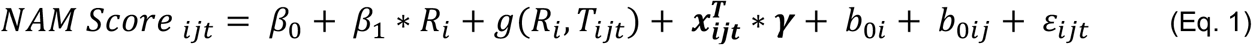

Where time function *g*(*R*_*i*_, *T*_*ijt*_ ):

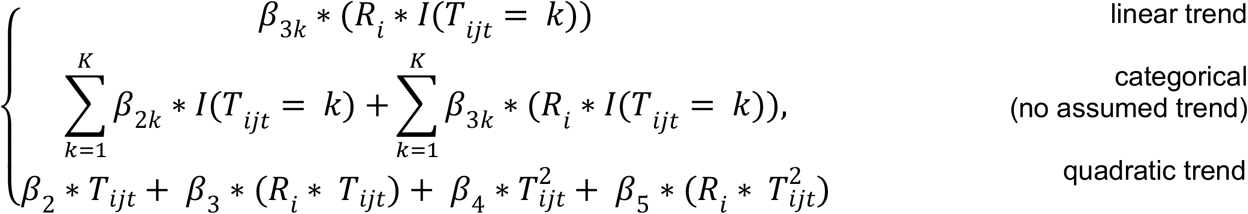

Here, i indexes subjects, j indexes cells and t indexes timepoints. **X**_**ijt**_ denotes the set of covariates included in the model. b_0i_ and b_0ij_ are subject-level and sample-level random intercepts respectively. K denotes the total number of timepoints, R_i_ indicates response status and T_ijt_ denotes time. The regression coefficients β correspond the fixed effect specified in the model.

#### Association score strategy prioritization

The approach for prioritizing NAM candidates differs based on the dimensionality reduction method generating the NAM scores. Because unsupervised/weakly supervised approaches (PCA or binary PLS-DA on the clinical outcome only) methods do not explicitly optimize for the time dependent clinical outcome effect during dimensionality reduction, we rank all components by LRT p-value to identify top candidates showing the strongest interaction effects. NAM scores are examined in order of statistical significance (LRT p-value), with lower p-values indicating stronger evidence for time dependent clinical phenotype patterns. In contrast, for PLS-DA with composite time-by-clinical outcomes (e.g., 6-level responder status at V0/V1/V2), the first latent component is explicitly optimized during dimensionality reduction to maximally discriminate between these groups. Because this component is constructed to capture time dependent outcome structure by design, LRT-based ranking provides limited additional information for candidate prioritization. Instead, the subsequent LMM analysis serves complementary purposes: providing formal statistical inference with p-values and confidence intervals, quantifying effect sizes in a standardized framework, adjusting for potential confounders, and characterizing between-subject heterogeneity through random effects variance components. In this supervised context, results can be used to guide biological interpretation.

### scLASER Simulation Method

*Simulation strategy*: We developed a generative simulator that produces longitudinal single-cell datasets with a user-defined sampling design, cell-type composition dynamics, and per-cell feature values. The simulator (i) creates study metadata (e.g. participants, visits/samples, covariates) alongside sample-specific cell-type compositions that can differ by group and time, and (ii) simulates per-cell feature matrices (either gene expression or low-dimension PCs) using a Gaussian Mixture Model. The model attributes variance to predefined factors, including cell-type clusters and sample-level categorical covariates, and exposes tunable parameters that directly control each factor’s contribution, allowing the proportion of variance explained (PVE) by each source to be specified.

*Flexible inputs for simulation*: We have designed and implemented key user-defined inputs in our scLASER simulator, including number of participants (n_individuals), visits (or samples) per participant (time_points), the number of cells per major cell type per sample (n_cells), group assignment specified by the proportion in the disease or treatment group (prop_disease), number of cell types (decomposed into n_major_cell_types and n_minor_cell_types), the subset of associated cell types (n_major_interact_celltypes and n_minor_interact_celltypes), optional covariates (i.e. batch, age, sex), the visit specific effect size (visit_effects_responder and visit_effects_nonresponder) which is the relative increase or decrease of the associated cell type proportion compared to the initial visit proportion for that specific group and the number of features (genes or PCs, n_features).

### Longitudinal Simulation Scenarios

Taking treatment response as a key clinical interest, individuals were monitored longitudinally to identify those who responded vs those who did not (non-responder). We simulated 10 cell types labeled A–J, including seven major (A–G) and three minor (H–J). Four cell types (A, B, I, J) were designated as associated with a response×time effect; the remaining types had no imposed group-by-visit signal. We simulated a variety of effect sizes ranging from 0.1 to 0.8, both expanded and depleted over time. The compositional variability parameter (sd_celltypes), which controls how much relative the per-sample fluctuation in cell-type counts fluctuate around the baseline cell count (n_cells) was fixed at 0.1 for all simulations.

Two-time-point design (V0, V1): for each associated cell type, we generated datasets with visit-specific effect sizes ranging from 0.1 to 0.8 (0.1, 0.2, 0.4, 0.6, 0.8). Cell types A and I were depleted over time in responders (negative trajectory), whereas B and J were expanded over time in responders (positive trajectory). We also varied participant number (n=10, 20, 30, 40, and 50 participants), number of cells per major cell type (n=100, 500 or 1,000), and sampling balance (either 1:1 or 2:1 responders:non-responders).

Three-time-point design (V0, V1, V2): we again simulated the 10 cell types (A–J) with the same four associated types (A, B, I, J). We had two different trends over time. First, we simulated linear trends where the associated cell types either expanded or depleted consistently, with per-visit increments between 0.05 and 0.40 (e.g., +0.05 at V1 and +0.10 at V2 relative to V0 for an expanding type). Secondly, we generated non-linear trends where the group difference peaked at V1 and attenuated at V2 (e.g., larger |effect| at V1 vs V0 than at V2 vs V0), capturing a bump pattern that is non-linear.

We also varied participant number, cells per sample, and sampling balance; memory constraints emerged as the primary limitation at large scales (>100 participants, >10,000 cells/sample). Comparison with interaction tests applied directly to cell-type frequencies confirmed that scLASER’s neighborhood-level analysis captures compositional structure beyond simple abundance shifts.

#### Evaluation of data reduction approaches

To inform the choice between PCA and PLS-DA for dimensionality reduction, we simulated a three–time-point longitudinal design (V0, V1, V2) with increased compositional variability relative to prior simulations. Specifically, the compositional variability parameter (sd_celltypes), controlling per-sample fluctuations in cell-type counts around the baseline cell count, was doubled from previous settings and fixed at 0.2 for all simulations. Each dataset consisted of eight major cell types, with no minor cell types included. Among the eight cell types, two were designated as associated with responder status. One cell type exhibited a main effect of responder status only, with a fixed effect size of 0.2 that was constant across visits. A second cell type exhibited a time-by-responder interaction, with visit-specific effect sizes of 0.0, 0.1, or 0.2 across V0–V2. The directions of effect were specified to be opposite between the two associated cell types, such that one cell type expanded over time in responders while the other depleted. All remaining cell types were simulated under the null, with no main or interaction effects.

#### Benchmarking performance and metrics

To benchmark scLASER against traditional frequency-based interaction tests, linear mixed effect models, we randomly generated 200 independent datasets for each temporal design. For 2-timepoint simulations, we set effect size to 0.2, 30 participants, and 100 cells per major cell type per sample. For 3-timepoint simulations, we applied non-linear effect sizes (0, 0.2, 0.1 at V0, V1, V2, respectively) with the same sample size parameters. Sensitivity was calculated from 2×2 contingency tables comparing true simulation status (associated vs. non-associated cell types) against detected status.

For scLASER, we first identified significant time-response NAM Scores (interaction LRT p-value < 0.05), then associated each significant NAM Score with cell-type abundances via linear mixed effects models (NAM score as outcome and cell type as predictor) with permutation testing (n = 100 permutations); cell types with a permuted p-value < 0.05 were classified as detected. This two-step approach mirrors real-world workflows where temporal signals are first identified, then interpreted.

As a benchmarking method, we compared scLASER’s performance against a frequency-based approach on the dataset we simulated Specifically, the frequency-based method we implemented is built based on generalized linear mixed effects models directly to test each cell-type cluster proportion with time-response interaction term incorporated in the full model (Eq 2). As model design, we have main effects for time and response status, specified covariates (age and sex), and random effects for samples nested within participants to account for within-subject correlation. In short, this serves as a benchmarking approach to evaluate the sensitivity of scLASER.

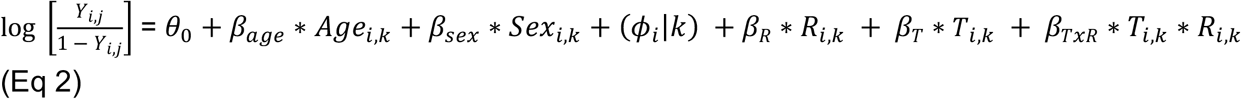

*Y*_*i,j*_ is the odds of cell *i* belonging to cluster *j, θ*_*j*_ is the intercept for cluster *j, β*_*age*_ and *β*_*sex*_ indicate the fixed-effect of age and sex for cell *i* from k^th^ sample, respectively; (*ϕ*_*i*_ |*k*) is the random effect for cell *i* from k^th^ sample, *β*_*R*_ indicates the effect of k^th^ sample’s response outcome status and variable *β*_*TxR*_ indicates the effect of k^th^ sample’s time dependent response outcome status for a specific cell i, at sample k. We compared the full model to a nested null model that excludes the time-response interaction term, but retains all main effects and covariates using a LRT. Cell types with an LRT p-value < 0.05 were classified as detected.

### Application on Real Longitudinal Datasets

#### Longitudinal IBD single-cell data for remission analysis

We applied scLASER to a longitudinal scRNA-seq dataset from Thomas et al.^4^. In brief, this study profiled scRNA-seq from five distinct gut regions in patients with inflammatory bowel disease (IBD), collecting samples both before and after treatment from 22 patients with ulcerative colitis (UC) and 16 patients with Crohn’s disease (CD). Because stromal cells are implicated in driving tissue inflammation and remodeling, we focused our analysis exclusively on this compartment(n = 103,503 cells).

Expression matrices were processed as in the original study, including quality control and normalization. Principal components were computed and harmonized using Harmony^15^ to correct for inter-individual and batch effects. The harmonized embeddings were used to construct a Neighborhood Abundance Matrix (NAM),, from which 30 NAM-derived principal components (NAM PCs) were retained and used in our longitudinal analysis pipeline. Specifically, we examined the interaction between treatment timepoint (pre-versus post-treatment) and clinical response (remission versus non-remission) as the primary effect of interest. Given only two timepoints, we treated time as a categorical variable (pre-vs post-). We included the following covariates: IBD subtype (UC or CD), biopsy location (terminal ileum, ascending colon, descending colon, sigmoid, or rectum), patient age, and gender. All available samples were retained in the analysis, including those without matched biopsy sites across timepoints.

#### Longitudinal COVID-19 single-cell data for disease severity analysis

The longitudinal COVID-19 single-cell dataset from Lin and Rajagopalan et al.^5^ profiled individuals with confirmed SARS-CoV-2 infection across multiple timepoints. We applied scLASER to this dataset to evaluate clinical trajectories of disease severity. Participants were classified as progressors (those whose disease severity worsened at any point after first visit) or non-progressors (those who maintained or improved in severity). Symptom duration was calculated as days from symptom onset, and visits were grouped into four temporal stages: days 1–4, 5–8, 9–14, and >14.Of the 108 total participants, 10 with only a single visit and 14 whose samples all occurred within the same temporal stage were excluded, as progression status could not be determined. The final analytic dataset included 84 participants (65 non-progressors and 19 progressors) with 238 individual samples.

From the processed single-cell dataset, we selected T and NK cells (188,181 cells) for downstream analysis. Expression matrices were preprocessed following the original study, including quality control,normalization, PCA, and Harmony-based correction for inter-individual and batch effects. The harmonized embeddings were used to construct a NAM summarizing local cell-state abundance across samples. 36 NAM-derived PCs were retained based on variance explained and used for all subsequent longitudinal analyses.

To capture temporal immune dynamics, we applied the scLASER pipeline to model each NAM PC as a function of time, disease progression status, and their interaction. Given the presence of more than two timepoints, time was modeled in three complementary ways: continuous, quadratic, and categorical (four levels), to assess both linear and nonlinear trajectories. For each NAM PC, the best-fitting model was selected based on AIC. All models adjusted for age, sex, and baseline disease severity.

## Research Standards

### Data Availability

The processed real-world single-cell datasets with metadata used in this study are publicly available through Zenodo. The IBD dataset can be accessed at https://doi.org/10.5281/zenodo.13768607, and the COVID-19 dataset can be accessed at https://doi.org/10.5281/zenodo.5153528.

### Code Availability

All code used for the scLASER framework, data processing, analysis, and simulation in this study is publicly available at https://github.com/fanzhanglab/scLASER.

## Acknowledgments

We thank the members of the Zhang lab for their insightful feedback, and we specifically thank Revanth Krishna for computational support.

## Contributions

Conceptualization: FZ, LAV

Methodology: LAV, JI, JV, CW

Software: JV

Simulation: JI

Formal analysis: LAV

Investigation: LAV

Validation: LAV, JV, JY

Visualization: LAV

Supervision: FZ

Writing—original draft: LAV, FZ

Writing—review & editing: LAV, FZ, JV, JI, JY, CW

