## Supplemental Figures for "scLASER: a robust framework for simulating and detecting time-dependent single-cell dynamics in longitudinal studies"

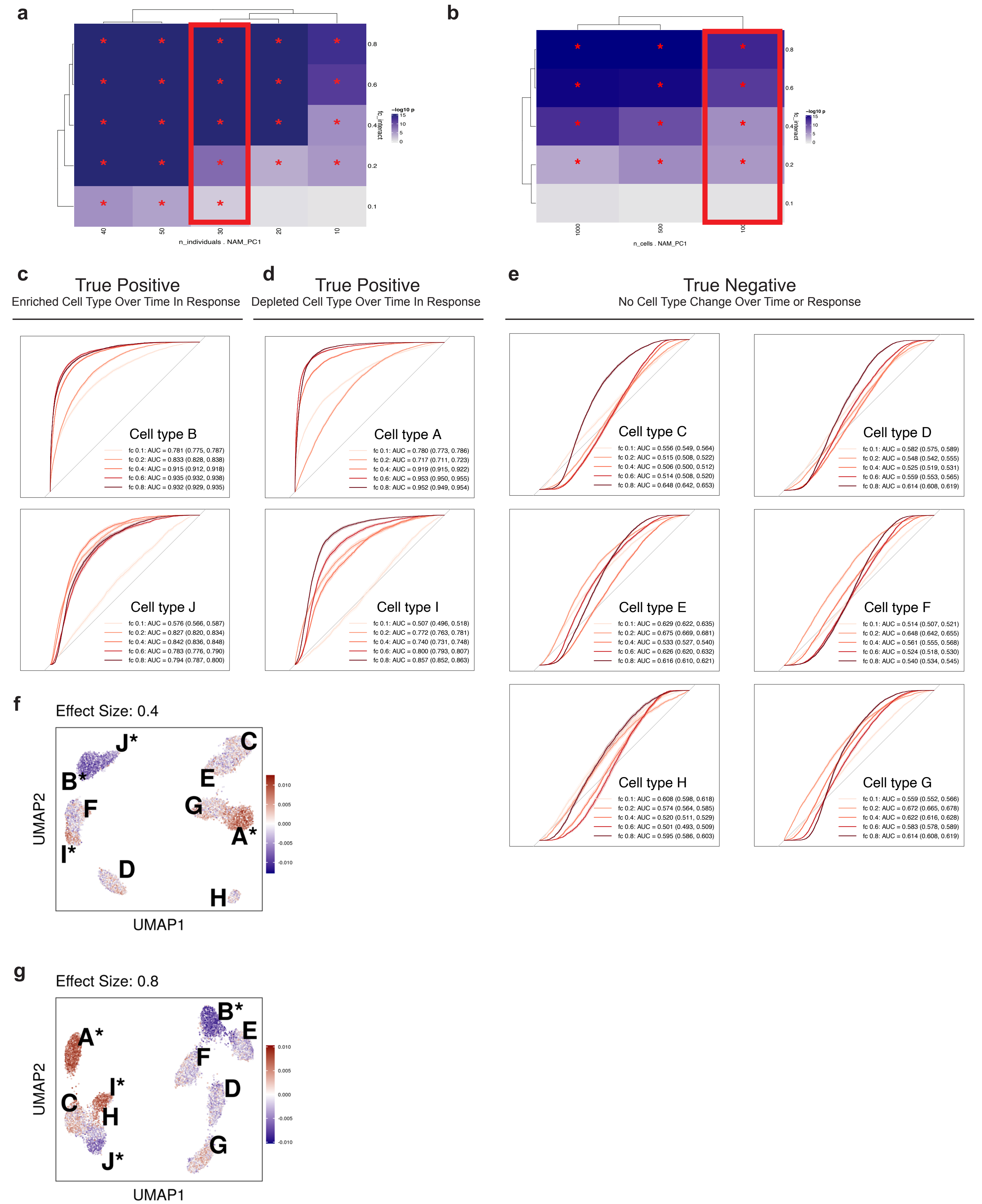


**Supplemental Figure 1. scLASER detection performance and ROC cell type classification across two-timepoint simulation parameters.**
(a) Heatmap showing -log10 p-values for the top NAM score (PC1) across varying numbers of individuals (10-50) and effect sizes (0.1-0.8), with fixed cell count of 100 per major cell type. *LRT p-value < 0.05. Red box highlights datasets used in subsequent ROC analyses and UMAP visualizations.
(b) Heatmap showing -log10 p-values for the top NAM score (PC1) across varying numbers of cells per sample per major cell type (100, 500, 1000) and effect sizes (0.1-0.8), with fixed sample size of 30 individuals. Red box highlights datasets used in subsequent ROC analyses and UMAP visualizations.
(c-e) ROC curves for all 10 simulated cell types across different effect sizes (fc 0.1-0.8) in 2-timepoint simulations. (c) True positive enriched cell types A and B showing high sensitivity with increasing effect sizes. (d) True positive depleted cell types I and J demonstrating detection of negative trajectories. (e) True negative cell types C-H maintaining specificity with AUCs near 0.5 across all effect sizes, confirming appropriate Type I error control.
(f-g) UMAP of simulated data colored by scLASER association scores (NAM PC1) for representative effect sizes. (f) Effect size 0.4.
(g) Effect size 0.8. Cell types A-J correspond to major A-G and rare H-J cell types; red indicates positive and blue indicates negative associations.

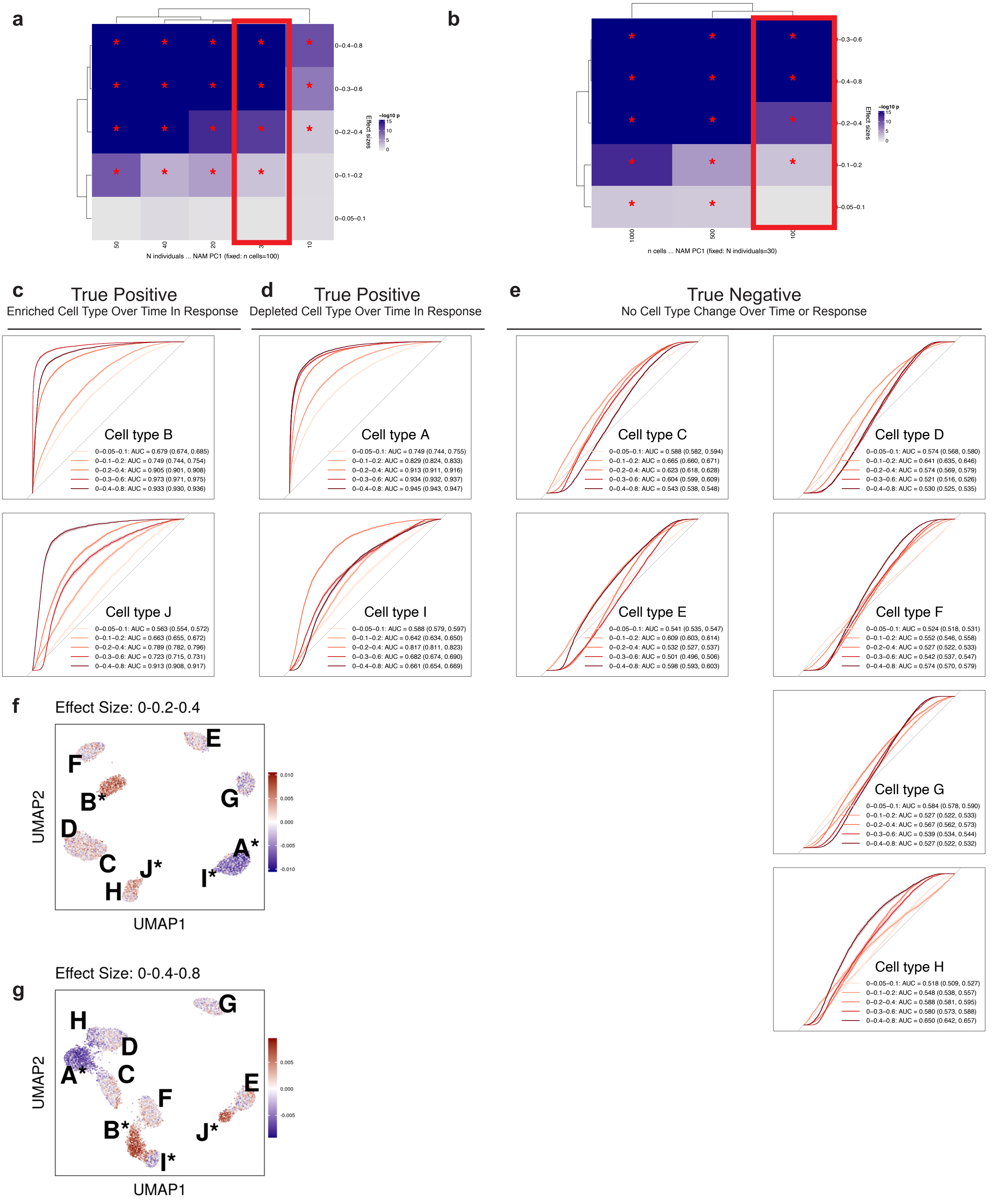


**Supplemental Figure 2. scLASER detection performance and ROC cell type classification across three-timepoint linear trend simulation parameters.** (a) Heatmap showing -log10 p-values for the top NAM score (PC1) across varying numbers of individuals (10-50) and linear trend effect sizes (0-0.05 0.1 to 0-0.4-0.8 representing incremental changes at V1 and V2 relative to V0), with fixed cell count of 100 per major cell type. *LRT p-value < 0.05. Red box highlights datasets used in subsequent ROC analyses and UMAP visualizations.
(b) Heatmap showing -log10 p-values for the top NAM score (PC1) across varying numbers of cells per sample per major cell type (100, 500, 1000) and linear trend effect sizes (0-0.05-0.1 to 0-0.4-0.8) with fixed sample size of 30 individuals. Red box highlights datasets used in subsequent ROC analyses and UMAPs.
(c-e) ROC curves for all 10 simulated cell types across different linear effect sizes in 3-timepoint simulations (V0, V1, V2). (c) True positive enriched cell types B and J showing sensitivity that increases with larger non-linear effect sizes in both major (B) and rare (J) cell types, demonstrating detection of monotonic expansion. (d) True positive depleted cell types A and I demonstrating detection of monotonic depletion in both major A and rare I cell types.
(e) True negative cell types C-H maintaining specificity with AUCs near 0.5 across all effect sizes, confirming appropriate Type I error control.
(f-g) UMAP visualizations of simulated data colored by scLASER association scores (NAM PC1) for representative non-linear effect sizes. (f) Effect size 0-0.2-0.4 (increments of 0.2 per visit). (g) Effect size 0-0.4-0.8 (increments of 0.4 per visit). Cell type A-J correspond to major A-G and rare H-J cell types; red indicates positive and blue indicates negative associations.

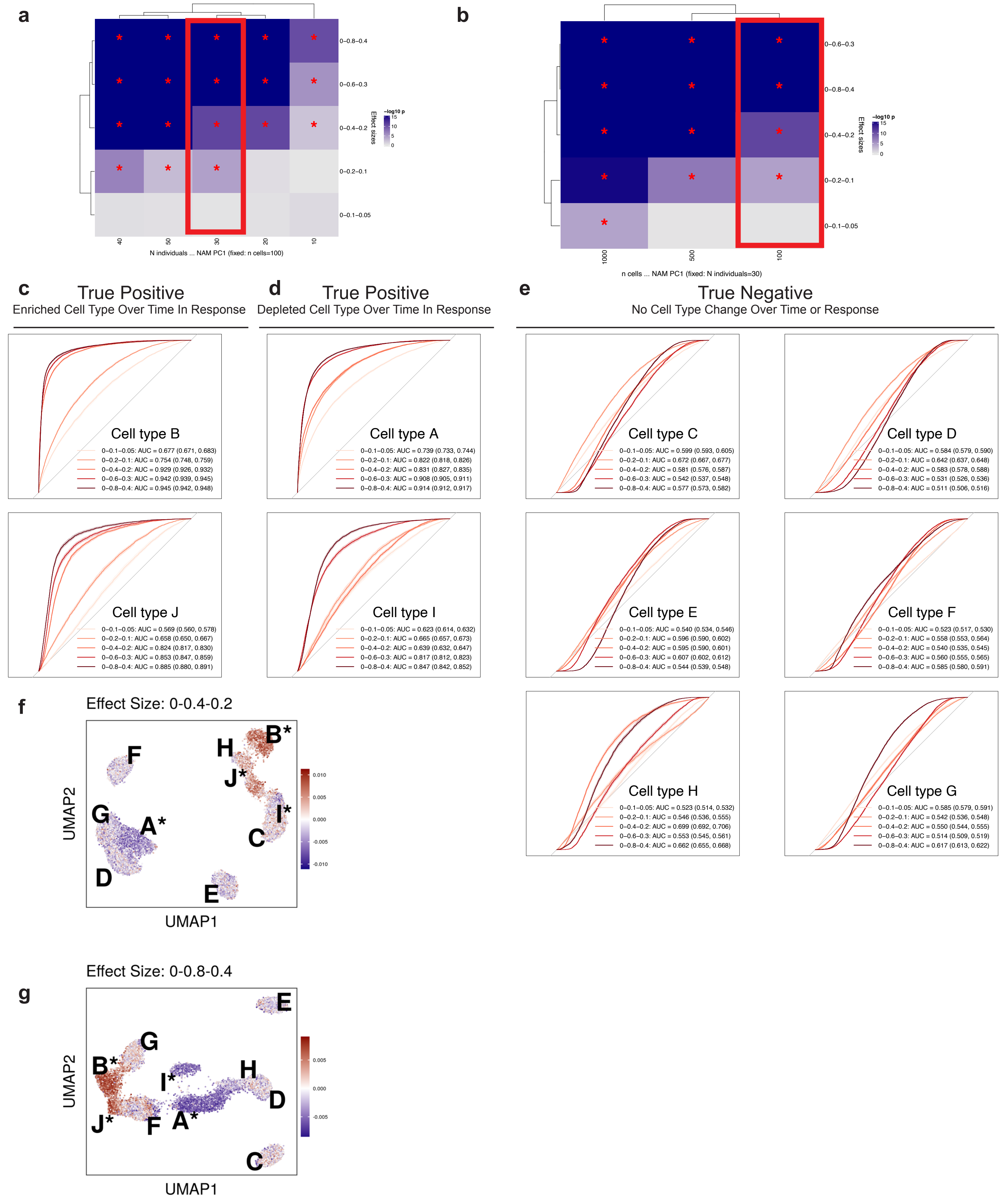


**Supplemental Figure 3. scLASER detection performance and ROC cell type classification across three-timepoint non-linear trend simulation parameters.** (a) Heatmap showing -log10 p-values for the top NAM score (PC1) across varying numbers of individuals (10-50) and non-linear effect sizes (0-0.1-0.05 to 0-0.8-0.4 representing a bump pattern where effects peak at V1 and attenuate at V2 relative to V0), with fixed cell count of 100 per major cell type. *LRT p-value < 0.05. Red box highlights datasets used in subsequent ROC analyses and UMAP visualizations.
(b) Heatmap showing -log10 p-values for the top NAM score (PC1) across varying numbers of cells per sample per major cell type (100, 500, 1000) and non-linear effect sizes (0-0.1-0.05 to 0-0.8-0.4) with fixed sample size of 30 individuals. Red box highlights datasets used in subsequent ROC analyses and UMAPs.
(c-e) ROC curves for all 10 simulated cell types across different linear effect sizes in 3-timepoint simulations (V0, V1, V2). (c) True positive enriched cell types B and J showing sensitivity that increases with larger non- linear effect sizes in both major B and rare J cell types, demonstrating detection of transient expansion patterns that peak at V1. (d) True positive depleted cell types A and I demonstrating detection of non-linear depletion trajectories in both major A and rare I cell types.
(e) True negative cell types C-H maintaining specificity with AUCs near 0.5 across all effect sizes, confirming appropriate Type I error control.
(f-g) UMAPs of simulated data colored by scLASER association scores (NAM PC1) for representative non-linear effect sizes. (f) Effect size 0-0.4-0.2 (peak effect of 0.4 at V1, attenuating to 0.2 at V2). (g) Effect size 0-0.8-0.4 (peak effect of 0.8 at V1, attenuating to 0.4 at V2). Cell types A-J correspond to major A-G and rare H-J cell types; red indicates positive and blue indicates negative associations.

**a**

**Frequency Cluster Based Testing Pipeline: 2 timepoint non-linear simulation**

All Cell Types

| | True Positive | True Negative |
|---|---|---|
| Significant | 755 | 0 |
| Non-significant | 45 | 1200 |

Rare Cell Types

| | True Positive | True Negative |
|---|---|---|
| Significant | 355 | 0 |
| Non-significant | 45 | 200 |

**b**

**scLASER Analysis Pipeline: 2 timepoint non-linear simulation**

All Cell Types

| | True Positive | True Negative |
|---|---|---|
| Significant | 1006 | 1382 |
| Non-significant | 26 | 166 |

Rare Cell Types

| | True Positive | True Negative |
|---|---|---|
| Significant | 499 | 220 |
| Non-significant | 17 | 38 |

**c**

**Frequency Cluster Based Testing Pipeline: 3 timepoint non-linear simulation**

All Cell Types

| | True Positive | True Negative |
|---|---|---|
| Significant | 247 | 1 |
| Non-significant | 553 | 1199 |

Rare Cell Types

| | True Positive | True Negative |
|---|---|---|
| Significant | 18 | 0 |
| Non-significant | 382 | 200 |

**d**

**scLASER Analysis Pipeline: 3 timepoint non-linear simulation**

All Cell Types

| | True Positive | True Negative |
|---|---|---|
| Significant | 994 | 1394 |
| Non-significant | 138 | 304 |

Rare Cell Types

| | True Positive | True Negative |
|---|---|---|
| Significant | 493 | 217 |
| Non-significant | 73 | 66 |

**Supplemental Figure 4. Benchmarking scLASER against frequency-based tests across simulation scenarios.**
(a) Contingency table summarizing results from 200 two-timepoint simulations (V0, V1) using frequency-based generalized linear mixed effects models (GLMM). Simulations used effect size 0.6, 30 individuals, and 100 cells per major cell type. True positives represent correct detection of cell types A, B, I, and J with time×response status effects. Overall sensitivity: 94.4% (rare cell types: 88.8%).
(b) Contingency table for scLASER across the same 200 two-timepoint simulations described in (a). scLASER demonstrates improved sensitivity, particularly for rare cell types. Overall sensitivity: 97.5% (rare cell types: 96.7%).
(c) Contingency table for frequency-based GLMM approach across 200 three-timepoint simulations (V0, V1, V2) with non-linear effect patterns where effect sizes progress from 0 at V0 to 0.2 at V1, then decrease to 0.1 at V2, maintaining 30 individuals and 100 cells per major cell type. Overall sensitivity: 30.9% (rare cell types: 4.5%).
(d) Contingency table for scLASER across the same 200 three-timepoint non-linear simulations described in (c). scLASER's adaptive model selection strategy substantially improves detection of non-linear temporal patterns. Overall sensitivity: 87.8% (rare cell types: 87.1%).

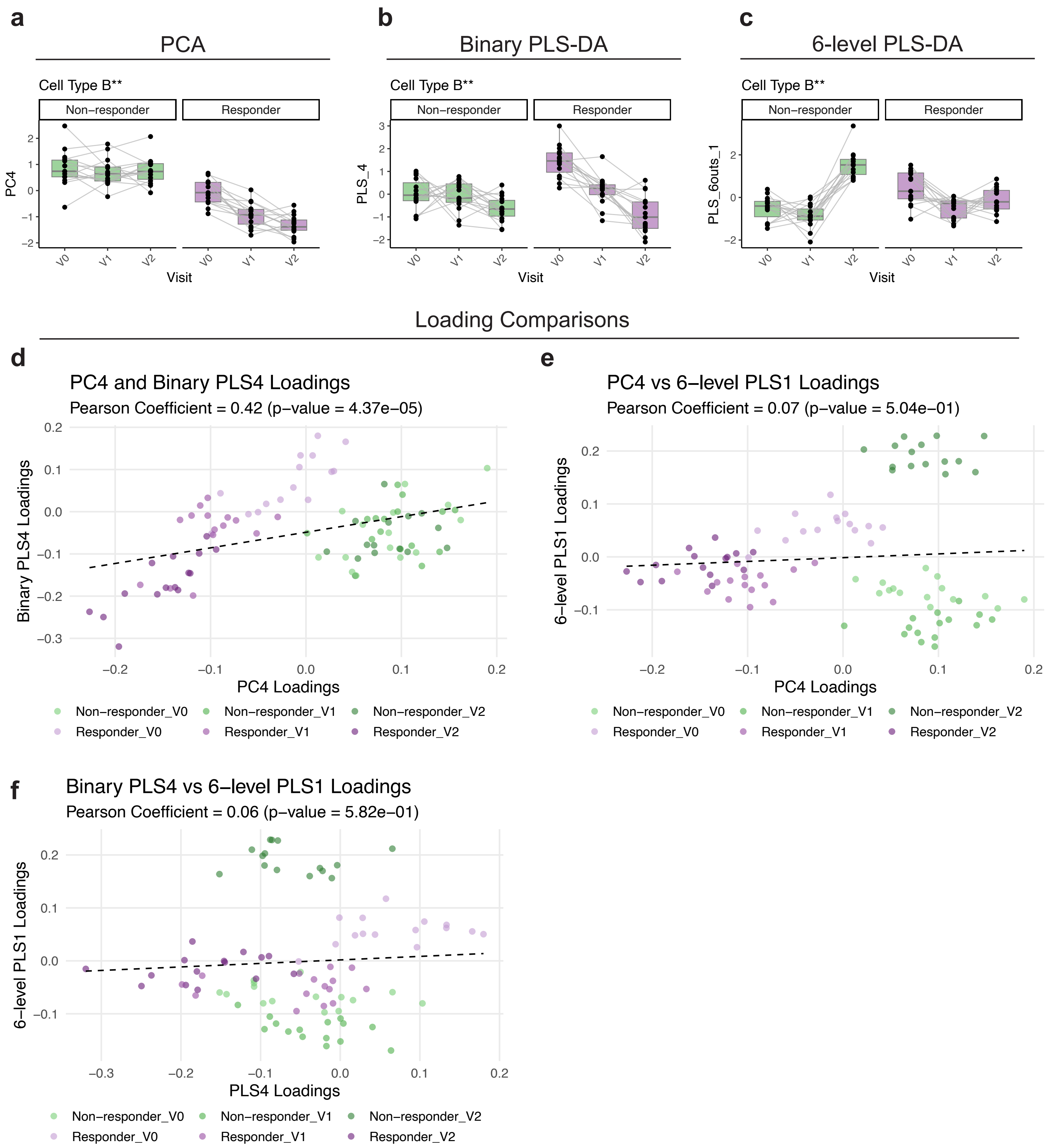


**Supplemental Figure 5. PCA and PLS-DA data reduction approaches yield distinct neighborhood loadings.**
a-c. Longitudinal association score trajectories within cell type B (simulated to have significant time-by-response status effect) for different data reduction techniques. (a) PC4 (top PCA-derived NAM score), (b) binary PLS4 (top binary response status PLS-DA NAM score), and (c) 6-level PLS1 (top 6-level time-by-response status PLS-DA NAM score). Individual data points represent samples, with lines connecting longitudinal measurements from the same individual.
d-f. Scatter plots comparing neighborhood loadings across data reduction techniques for the top NAM score from each approach. Pearson correlation coefficients are reported with p-values. Neighborhoods are color-coded by response status (responder: purple; non-responder: green) and time (earlier visits: lighter shades; later visits: darker shades). (d) PC4 vs binary PLS4 loadings (r=0.42, p=4.37e-05), (e) PC4 vs 6-level PLS1 loadings (r=0.07, p=0.504), (f) binary PLS4 vs 6-level PLS1 loadings (r=0.06, p=0.582). Low correlations between PCA and 6-level PLS-DA approaches demonstrate that different reduction methods capture distinct aspects of neighborhood structure.

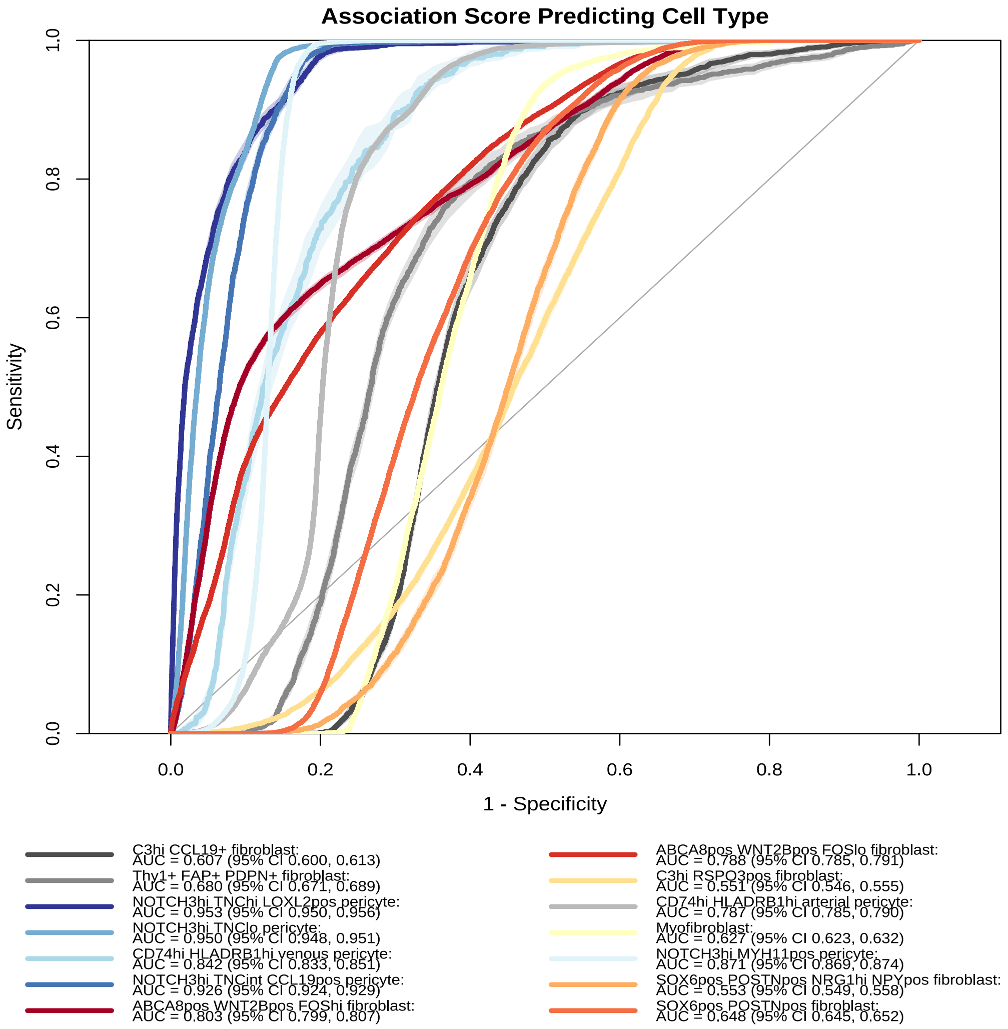


**Supplemental Figure 6. scLASER identifies a strong remission and post-treatment–specific effect.**
ROC curves of association score predicting all 14 stromal cell types. AUC (95% CI) are reported for each curve.
